# ZBTB7B is a Master Regulator of Salivary Acinar Cell Function

**DOI:** 10.64898/2026.10.09.758069

**Authors:** Thomas J. F. Pranzatelli, Paola Perez, Drew G. Michael, Kalie Dominick, Melanie Vacchio, Changyu Zheng, Daniel Martin, NIDCR/NIDCD Genomics and Computational Biology Core, Philip L. F. Johnson, Blake M. Warner, John A. Chiorini

## Abstract

Salivary glands exhibit transcriptional programs that distinguish them from other tissues, yet the upstream regulatory logic underlying these programs is not well defined. To identify transcription factors (TFs) that structure salivary identity, we generated RNA-seq and ATAC-seq from human major and minor salivary glands and inferred TF→TF regulatory edges from high-confidence chromatin footprints. ZBTB7B emerged as the most connected regulator across salivary samples. To locate regulatory regions associated with salivary-specific transcription, we trained hidden Markov models (HMMs) on chromatin accessibility across 4 Mbp windows surrounding salivary-specific and non-salivary-specific transcription start sites. The resulting chromatin states distinguished promoter-proximal openness from distal enhancer activity and identified salivary-specific enhancers enriched for ZBTB7B motifs. These enhancer states correlated strongly with expression of salivary genes, including AQP5, indicating that distal regulation contributes substantially to salivary transcriptional specificity.

To assess the consequences of losing Zbtb7b, we performed single-cell RNA-seq and spatial transcriptomics on submandibular and parotid glands from Zbtb7b knockout mice. Knockout mice exhibited reduced salivary flow, shifts in acinar cell proportions, and the emergence of a neonate-like acinar population characterized by loss of canonical secreted proteins and increased expression of MHC class I genes. These transcriptional and cellular changes occurred despite preserved gland architecture, suggesting that Zbtb7b stabilizes adult acinar identity rather than supporting structural maintenance. Together, these data identify ZBTB7B as a master regulator of salivary transcriptional networks, acting through both promoter-proximal and enhancer-level mechanisms, and demonstrate that loss of Zbtb7b disrupts acinar identity and impairs salivary function.

## Introduction

There are four kinds of salivary glands: three major salivary glands, the parotid, submandibular and sublingual; alongside labial or minor salivary glands. These glands perform different functions in the mouth. The submandibular glands, for example, secrete mucins that moisten the mouth and dampen bacterial growth, while the parotid glands are responsible for digestive secretions in response to consuming food. All of these glands are complex branching tissues that produce grams of protein and move large amounts of water into the mouth. Loss of the function of these glands leads to dryness, significant oral pain, and dental caries.

While the functional differences between the glands are well-characterized, the gene expression differences driving these functional changes (beyond the expression of specific secreted factors) are less well-understood. Upstream of these gene expression differences are the genes that regulate expression, the transcription factors (TFs). TFs bind in complexes around transcription start sites and recruit or prevent the recruitment of RNA polymerase. TF binding events will block DNA endonuclease activity and can be detected, like nucleosomes, with assays of chromatin accessibility. The low binding kinetics of TFs makes it possible to distinguish binding events from nucleosome complex occupancy and the signatures of TF binding are called footprints. When TFs bind to a piece of chromatin an educated guess can be made at the identity of the TF based on the sequence of DNA the TF binds to.

TFs that regulate multiple other TFs and, through them, can drive a cellular function are called “master regulators.” Examples of master regulators and their regulated function are MYBL1 and male meiosis(1), Snail and the epithelial-mesenchymal transition(2), and KLF5 in controlling the tumor microenvironment(3). Likely the most famous master regulators are the Yamanaka factors Oct4, Sox2 and Nanog, which, alone or in tandem with Gabpa, can drive the acquisition of pluripotency(4, 5). Sox2 is necessary for the differentiation of acinar cells in the developing salivary gland and is present in all epithelial cell types in adult salivary gland(6). In salivary glands, Sox2 directly affects expression of both terminal acinar markers like the water channel Aqp5 and acinar-specific TFs like Mist1, making it a candidate for a master regulator of salivary acinar function.

In this paper, RNA- and ATAC-seq were generated on major and minor salivary glands from human biopsies and autopsies. The TF ZBTB7B (Th-POK, c-Krox) was found consistently regulating other TFs across all glands sampled. ZBTB7B performs different roles in different tissues. Under the name Th-POK, ZBTB7B is most well-known as a TF necessary for CD4+ T cell activation(7–10). Through its expression in CD4+ T cells, ZBTB7B is a central part of the response to colorectal cancer(11), cerebral malaria(12) and radiation-induced aseptic inflammation(13). Particularly high levels of ZBTB7B in CD4+ T cells are found in HIV “elite controllers,” who are HIV-positive but maintain low viral loads(14). Less well-known, Th-POK is also involved in the differentiation of the myeloid lineage(15) (macrophages, monocytes, dendritic cells and granulocytes) and high expression of Th-POK in marrow macrophages inhibits osteoclast formation(16). Under its oldest name, c-Krox, ZBTB7B/Zbtb7b has been known to positively regulate the deposition of type I and type II collagen(17–19). It has previously been identified as a key regulator of mammary tissue: Zbtb7b knockout mice do not lactate(20) and ZBTB7B has been shown to canalize breast cancer cells to an epithelial identity(21). The gene is a central player in the progression of steatohepatitis(22) and hepatocellular carcinoma(23, 24). Zbtb7b drives pit cell differentiation in gastric organoids(25) and loss of ZBTB7B in gastric cancer is associated with immune escape(26) and increase in disease severity(27). Mutations and changes in ZBTB7B expression within tumors have been linked to disease severity in lung adenocarcinoma(28), pancreatic cancer(29), epithelial ovarian cancer(30) and bladder cancer(31). Through RSK3, ZBTB7B regulates ATXN1 in spinocerebellar ataxia type 1(32) and is linked to neural tube defects following treatment with benzo[a]pyrene(33). Zbtb7b binds to the lncRNA Blnc1 and downregulates the adipokine sST2 to promote white-to-brown fat differentiation(34, 35) and a variant in ZBTB7B has been linked to obesity in humans(36). Zbtb7b can be used to induce a primed-to-naive transition in embryonic stem cells, including expression of Yamanaka factors and other major zygotic genome activation genes(37). The goal of this paper is to clarify the role of ZBTB7B, the set of TFs regulated by ZBTB7B, and downstream genes of ZBTB7B in adult salivary glands, and the effects of losing this multi-functional transcription factor on the function of those glands.

To better understand the role of ZBTB7B in salivary glands, bulk RNAsequencing/ATAC-sequencing, single-cell RNA-seq, and Xenium spatial transcriptomics was performed on a transgenic mouse line of Zbtb7b knockout mice. Both parotid and submandibular glands were taken from these mice and analyzed separately. We report changes in transcription and proportion of cell types in submandibular glands in knockout mice. We also report the emergence of an unusual epithelial cell type specific to the parotid glands of knockout mice and show that these cells are segregated spatially within the gland. These mice lose salivary flow, suggesting that shifts in transcription, cellular composition of the gland, and cell identity correspond with reduced salivary function. These data point towards a role for ZBTB7B in late salivary development and as a master regulator of adult salivary function.

## Methods

### Human Patients

Human major salivary gland samples were acquired from the Human Cooperative Tissue Network. Human minor salivary gland samples were taken from healthy volunteers at the NIDCR Dental Clinic. Sections of tissue were immersed in 10% formalin and embedded in paraffin. Tissues were dissected by hand with microtome blades and then digested with liberase in a GentleMACS Octo for 45 minutes.

### RNA-sequencing

Samples were further disrupted with a handheld Qiagen TissueRupter and cells were lysed using QiaShredder tubes (Qiagen). RNA was isolated with an RNeasy Kit (Qiagen) and DNA digested with Turbo DNase (ThermoFisher). Samples were sequenced on an Illumina HiSeq at 150 x 150 bp for a total of 60 million reads per sample.

### ATAC-sequencing

Nuclei were isolated from salivary gland cell mixtures using IGEPAL-CA630 following the original protocol(38). Samples were indexed following the original protocol and quality assessed with qRT-PCR. All samples had a quality score of 6 or lower, suggesting high library diversity. Samples were sequenced on an Illumina HiSeq at 37 x 37 bp for a total of 200 million reads per sample.

### ATAC- and RNA-seq Analysis

ATAC-seq samples were processed and analyzed using the ATAC2GRN pipeline, producing a gene regulatory network(39). RNA-seq was quantified using Salmon in mapping-based mode with the --validateMappings flag(40, 41) and TFs binding sites were identified using CisBP(42) and mapped to the genome using FIMO(43). Only transcription factors expressed at a level of 0.5 CPM were included as possible regulators in a sample. With TF binding sites of different lengths and with A/Ts more common in the genome than G/Cs, each TF binding site motif has a different distribution of possible scores, making it necessary to compare individual TF binding sites as percentiles within their TF binding score distributions. Samples were said to have a TF regulating a gene if there was a footprint within the 5kb upstream and downstream of that gene’s transcription start site (TSS) overlapping a binding site for that TF which matched the genomic sequence in that interval with a score in the 95th percentile of all matching binding sites for that TF across the genome. When multiple TFs had binding sites overlapping a footprint, the TF with the highest percentile was chosen. The TF connection map was pruned to only include TFs that regulated at least one gene, and then pruned to only include TFs that regulate other TFs.

### Hidden Markov model (HMM)

Transcripts were grouped by transcription start sites (TSS) and total transcription from each TSS was summed. 1000 TSSs specific to salivary glands and 1000 TSSs non-specific to salivary glands were identified by sorting Mann-Whitney U statistics of rank difference between salivary and non-salivary samples. A hidden Markov model with 25 hidden states was trained on chromatin accessibility in a 2 Mbp window around each of these 2000 TSSs using a Poisson distribution for emission and a simple triangular rolling window of 100bp. Lambda estimates of the mean emission for each sample were calculated for each state and used to estimate a manifold(44). HMM states were clustered using Leiden clustering(45) and correlation measured between the mean estimate of accessibility and expression of that TSS.

### Mice

ZBTB7B−/− and +/−mice were acquired from the NCI and added to NIDCR protocol 21-1057-A2. Wild-type C57BL/6J mice were purchased from JAX. Salivary flow was measured via cannulation and submandibular and parotid glands were dissected from these mice. Sections of each gland were fixed in formalin and embedding in paraffin and the remainder were digested in liberase and processed for scRNA-seq libraries on the 10X Chromium machine. Formalin-fixed paraffin-embedded tissues were used with the Xenium kit (10X) to produce spatial maps of expression. Both publicly available and newly generated libraries were processed using scanpy(46).

## Results

### ZBTB7B is a master regulator of transcription factors

Major and minor salivary glands share a pattern of expression that distinguishes them from non-salivary tissues, particularly expression of transcription factors (TFs; Figure 1a, Supplementary Figure 1a). This pattern of transcription discriminates salivary glands from other tissues both via clustering and dimensionality reduction, and further discriminates diseased salivary glands from healthy glands (Figure 1b, Supplementary Figure 1b). Chromatin accessibility around transcription start sites is insufficient to distinguish salivary from non-salivary tissues (Supplementary Figure 1c), suggesting an additional factor driving the pattern of expression characteristic of salivary glands. When comparing connections between TFs, where one TF regulates another, and limiting analysis to those TF −> TF connections shared across salivary gland samples, ZBTB7B, expressed in major and minor salivary glands in males and females, was identified as a “master regulator” of other TFs, regulating by far the greatest number of other TFs in salivary glands (Figure 1c, Supplementary Figure 1d.)

**Figure 1.**
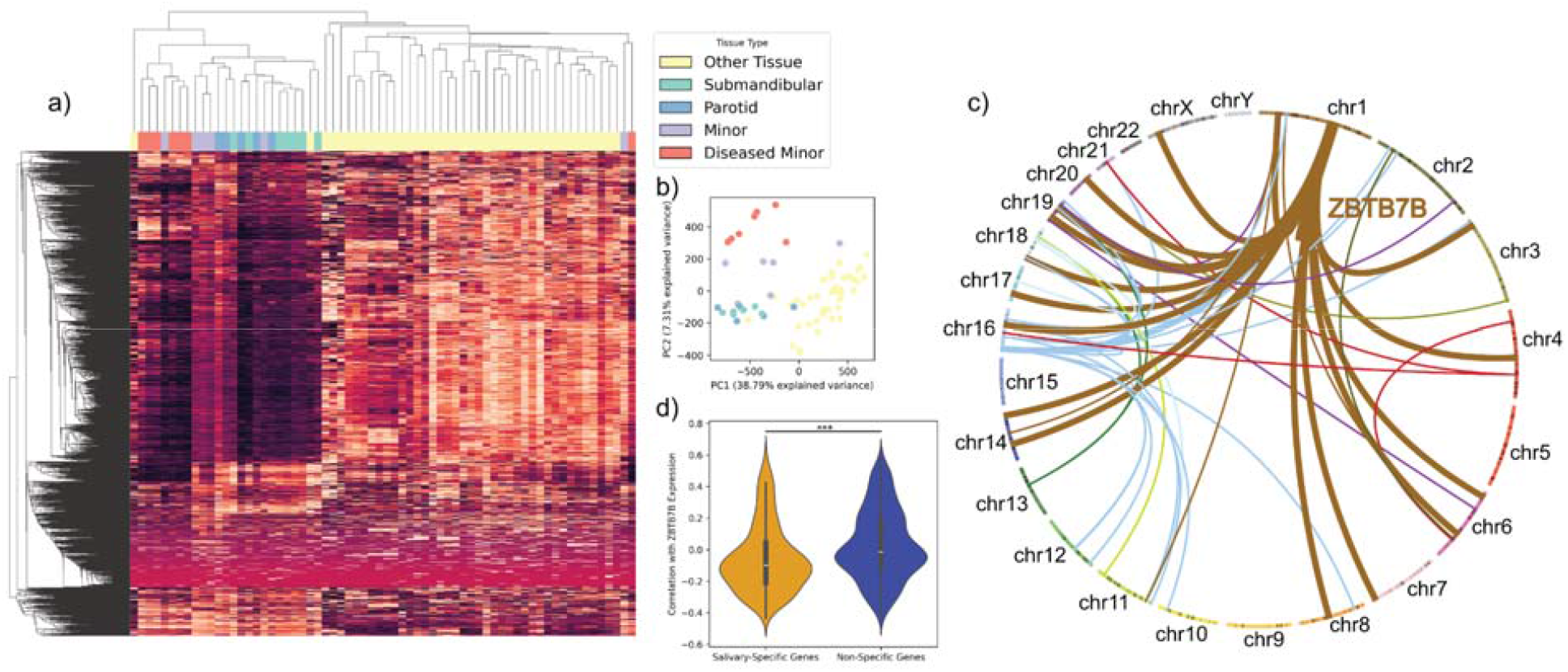
ZBTB7B is a master regulator of other transcription factors in adult human salivary glands. (a) Salivary and ENCODE RNA-seq samples (columns) clustered by expression of TFs (rows). Healthy salivary glands share a pattern of TF expression. (b) PCA plot of salivary and ENCODE tissue samples, based on TF expression. Healthy salivary glands cluster together, away from diseased and non-salivary tissues. (c) Circos plot linking TFs expressed in human salivary glands to binding events across the genome, colored by the chromosome where the regulating TF’s gene body is found. The TF regulating the greatest number of other TFs is ZBTB7B (shown with thicker brown lines). (d) Expression of ZBTB7B correlates with non-salivary-specific genes both positively and negatively, suggesting dual roles as an activator and suppressor. ZBTB7B has a higher chance of correlating negatively with salivary-specific genes.

Transcripts were grouped by TSS and the sum expression coming off of a TSS was compared as ranks between salivary and non-salivary tissues using Mann-Whitney U statistics. These most significant rank differences were used to identify 1000 TSSs specific to salivary glands, and 1000 TSSs specific to non-salivary tissues (“non-salivary-specific”). Salivary- and non-salivary-genes correlated with ZBTB7B expression both positively and negatively, suggesting dual roles as an activator and repressor (Figure 1d). Salivary-specific genes have a statistically significantly higher chance of correlating negatively with ZBTB7B.

### Hidden Markov model locates salivary-specific enhancer regions enriched for ZBTB7B motifs

Hidden Markov models (HMMs) with up to 25 hidden states were trained on 4 Mbp loci centered on the top 1000 salivary- and non-salivary-specific TSSs. These hidden states correspond to genomic intervals that share a pattern of chromatin accessibility in salivary and non-salivary samples. Across loci, constitutive and facultative heterochromatin could be distinguished from intergenic euchromatin, promoter regions and salivary-specific enhancers using their shared patterns of accessibility (Figure 2a, Supplementary Figure 2b,d). Openness in promoter elements correlates positively with salivary and non-salivary transcription while openness in cis-regulatory elements located farther from the TSS has greater variance in and higher average correlation with salivary expression (Figure 2b). These distal regulators contain higher numbers of ZBTB7B motifs.

**Figure 2.**
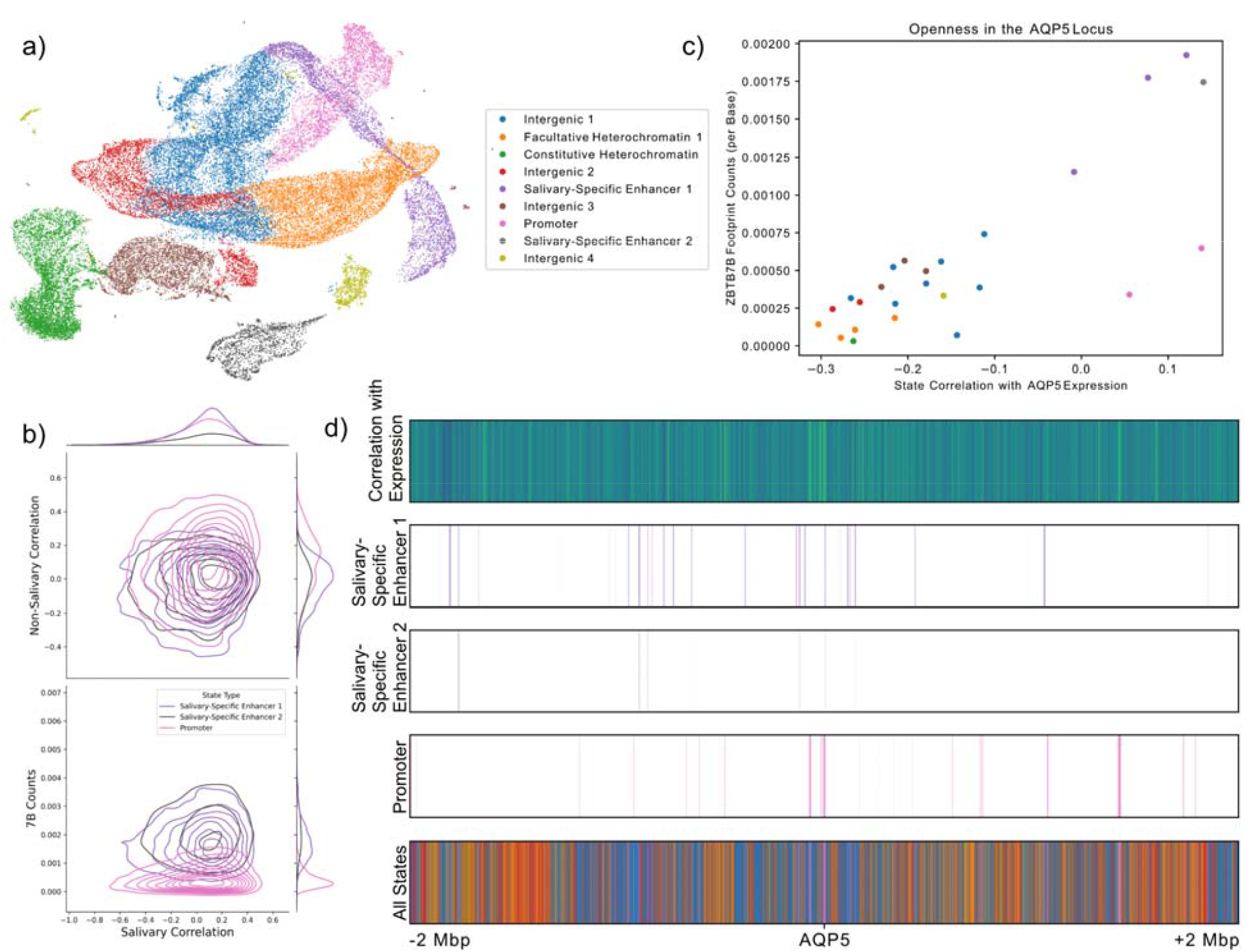
ZBTB7B motifs are found in salivary-specific enhancers. (a) UMAP of HMM states across 2000 salivary-specific and non-salivary-specific genes. States are Leiden clustered by pattern of ATAC openness across samples. (b) Correlation of openness with expression and average ZBTB7B motif counts per base are plotted for promoters are enhancer states. All three state types have positive correlation with salivary expression. Non-specific promoters have higher correlation with non-salivary samples, while salivary-specific cis-regulators have higher motif counts. (c) Correlation and motif counts are plotted for the twenty-five states in the 4 Mbp around AQP5 and colored by state type. Salivary-specific enhancers contain ZBTB7B motifs and correlate with AQP5 expression. (d) Genomic regions corresponding to each state in the AQP5 locus are plotted alongside correlation at that genomic coordinate and expression of AQP5.

AQP5 is an example of a salivary-specific gene that is positively correlated with openness in two promoter states and four salivary-specific enhancer states (Figure 2c). Multi-kilobase regions of positive association with expression were annotated by the HMM as promoter-like based on their pattern of accessibility, whereas shorter regions more distal from the TSS that nonetheless correlate with expression were classified as salivary-specific enhancers (Figure 2d).

### Newborn submandibular gland acini can be divided into multiple groups, which mirror groups seen in adult submandibular glands

To better understand the role of Zbtb7b in mouse salivary gland development, publicly available dissected embryonic mouse salivary gland single-cell RNA-seq was analyzed. Cells from developing submandibular glands were ordered by age and expression of key acinar cell genes plotted (Supplementary Figure 3a,b,d). Zbtb7b expression increases with development time and peaks at birth (Supplementary Figure 3c). This pattern is shared by other key salivary gland genes, including Wfdc12, Wfdc18 and Kit. Dcpp1 and 2, both regulated by Zbtb7b, also increase with developmental time. Dcpp1’s ortholog, ZG16B, is expressed most highly by PRR4+CST3+WFDC2-seromucous cells in adult humans. Amylase is consistently low in the submandibular gland and is a marker of parotid-specific function. Submandibular gland cells of newborn mice have acinar epithelium with broadly three patterns of expression: a ductal-like expression pattern that contains ductal markers like Wfdc18 and Krt8 but no Egf and instead Cldn3 (Set 1); an acinar subtype that produces parotid-like protein secretions including Bpifa2 (Set 2), and a majority population that expresses secreted mucins and canonical submandibular markers like Smgc (Set 3; Figure 3a-c, Supplementary Figure 3e). In addition to high levels of secretory activity, Set 2 most highly expressed Zbtb7b and Dcpp1 and 2. The expression patterns of Set 2 and Set 3 differ markedly in terms of proteins secreted but both express water-moving aquaporing Aqp5.

**Figure 3.**
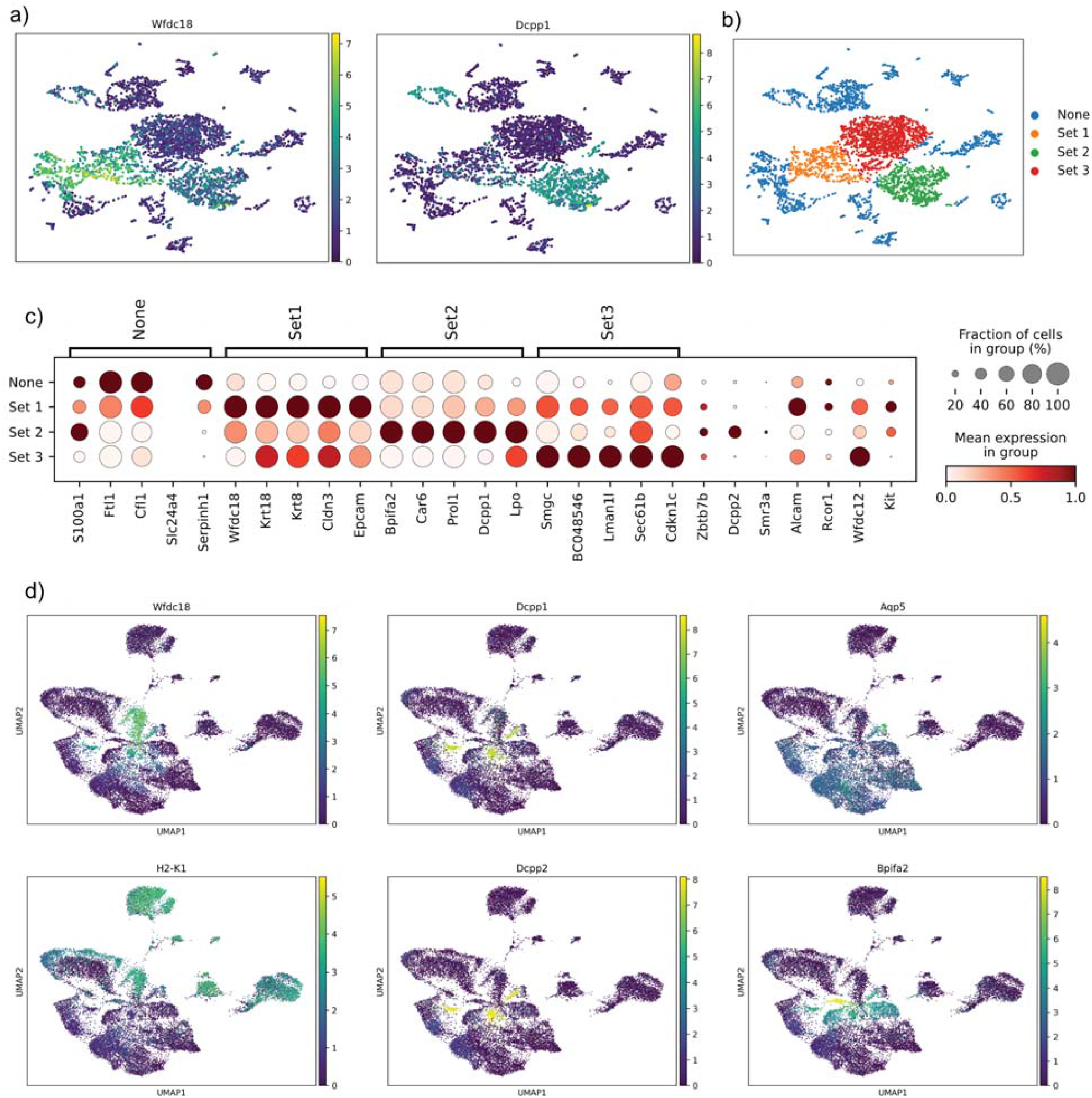
Developing mouse submandibular glands contain three types of acinar cells, which can be seen in the adult mouse. (a) Expression of Wfdc18 and Dcpp1 in salivary epithelial cells in newborn mouse submandibular. (b) Salivary epithelium in submandibular glands of newborn mice can be subdivided into three groups: an intercalated ductal-like Wfdc18 population (Set 1), a Dcpp1+ population of acinar cells (Set 2) and a third, majority population of acinar cells (Set 3). (c) Dotplot of gene expression specific to each subset of cells, normalized per-gene. Zbtb7b is least highly expressed in the Wfdc18+ population of acinar cells. (d) Feature plots of gene expression in the submandibular glands of Zbtb7b knockout and wild-type adult mice. Three cell types with three distinct patterns of expression – a Wfdc18+H2-K1+, a Dcpp1+Dcpp2+ and a Bpifa2+ cell type – are visible near the center of the UMAP.

Submandibular glands from homozygous knockout mice were not clearly different structurally from heterozygous or wild-type mice and all three had acinar structures producing mucins. To interrogate the identity of this new minority population, single-cell RNA-seq data was generated on all three genotypes of mice of multiple ages and sexes. A uniform manifold (UMAP) was built on K-nearest neighbors with batch balancing and projected into two dimensions (Figure 4d).

**Figure 4.**
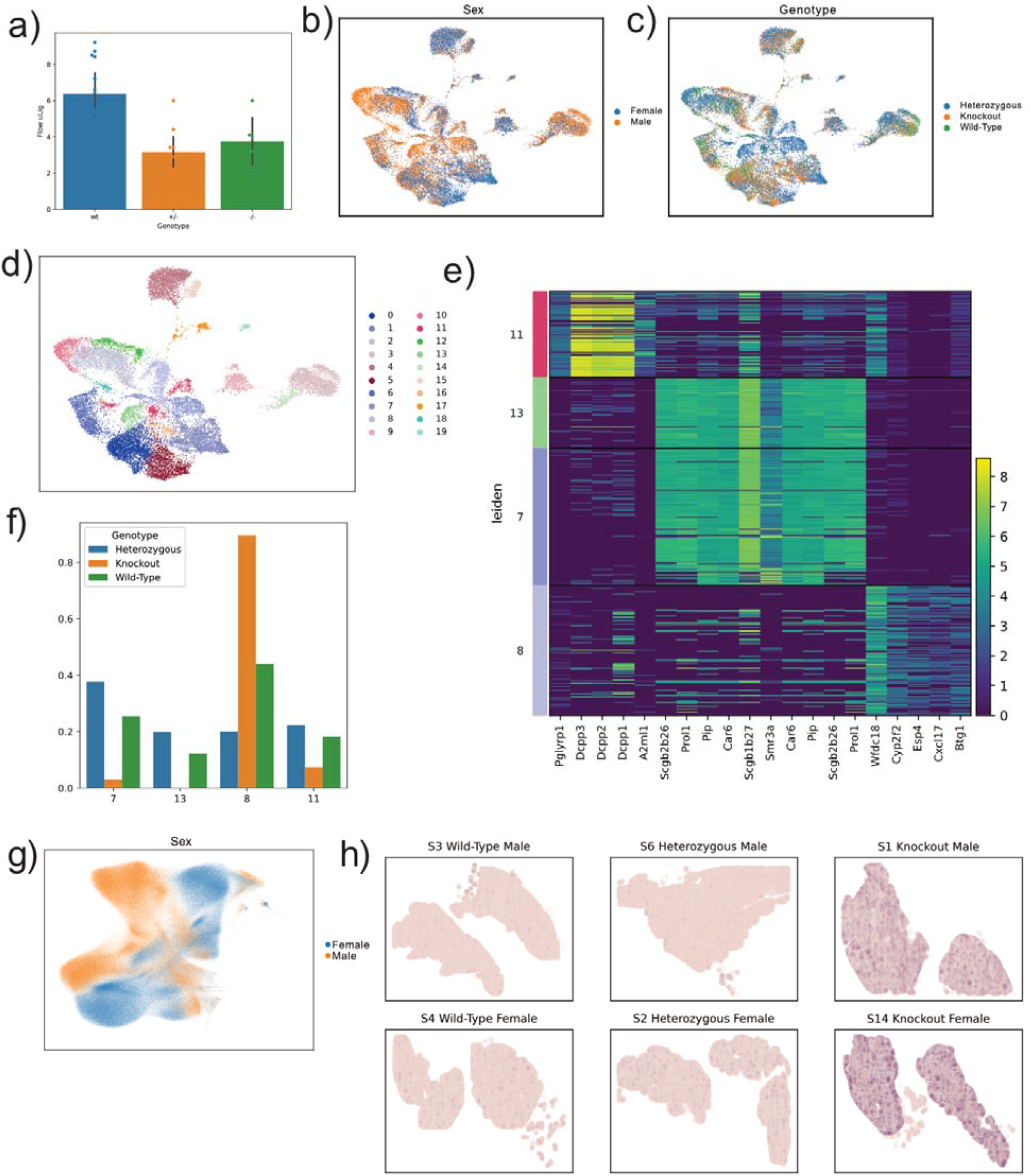
Heterozygous and wild-type mice have different ratios of acinar cells in their submandibular glands and salivate less than wild-type mice. (a) Both mice heterozygous and homozygous for Zbtb7b knockout have significantly reduced stimulated salivary flow. (b) UMAP of submandibular gland cells colored by sex. (c) UMAP of submandibular gland cells colored by genotype. (d) Leiden clustering of the submandibular gland. The three acinar expression patterns correspond to clusters 7 and 13, cluster 8 and cluster 11. (e) Proportion of cells found in each of the four clusters, limited to cells in those clusters. The vast majority of knockout cells are found in H2-K1+ Dcpp1-low cluster 8. (f) Heatmap of gene expression across the four clusters. Cluster 8 expresses a unique set of secreted proteins instead of the standard secreted proteins. (g) UMAP of Xenium quantification of submandibular glands. Differences are dominated by sex dimorphism. (h) Xenium quantification of Dio1 expression in the submandibular glands of male and female knockout and wild-type mice.

Three groups of acinar cells, similar to the groups found in developing submandibular, could be distinguished (Figure 3d, Supplementary Figure 3f). Set 1-like cells (Wfdc18+, “Cluster 8”) expressed high levels of H2-K1, B2m, Cd74 and Ano1, and low levels of Aqp5 and Kcnn4, and were highly enriched in mice homozygous for a Zbtb7b knockout (Figure 4c,f). In both the single-cell and Xenium data, the greatest differences between samples were driven by sexual dimorphism between male and female submandibular glands (Figure 4g, Supplementary Figure 4e). The proportions of these cell types, which changed with genotype, included changes in transcription of few genes including Dio1, and these were visible as changes occurring evenly across the gland (Figure 4h, Supplementary Figure 4a,b,f,g).

### Functional loss in salivary glands of Zbtb7b knockout mice

Salivary glands were cannulated and salivary flow was measured in wild-type mice and mice heterozygous and homozygous for Zbtb7b knockout. Mice with even one copy of knockout had statistically significantly less salivary gland flow (Figure 4a). No statistically significant difference was seen between heterozygotes and homozygotes.

### New neonate-like acinar cell type found spatially segregated in parotid mouse glands lacking Zbtb7b

Similar to submandibular glands of knockout mice, homozygous knockout parotid glands contained an enrichment for a subset of acinar cells (Figure 5a,f). Unlike in the submandibular gland, these cells have unique patterns of expression not found in heterozygous and wild-type tissue. These acinar cells lack alpha amylase (a marker of parotid function) and express Alcam, normally a ductal marker (Figure 5b).

**Figure 5.**
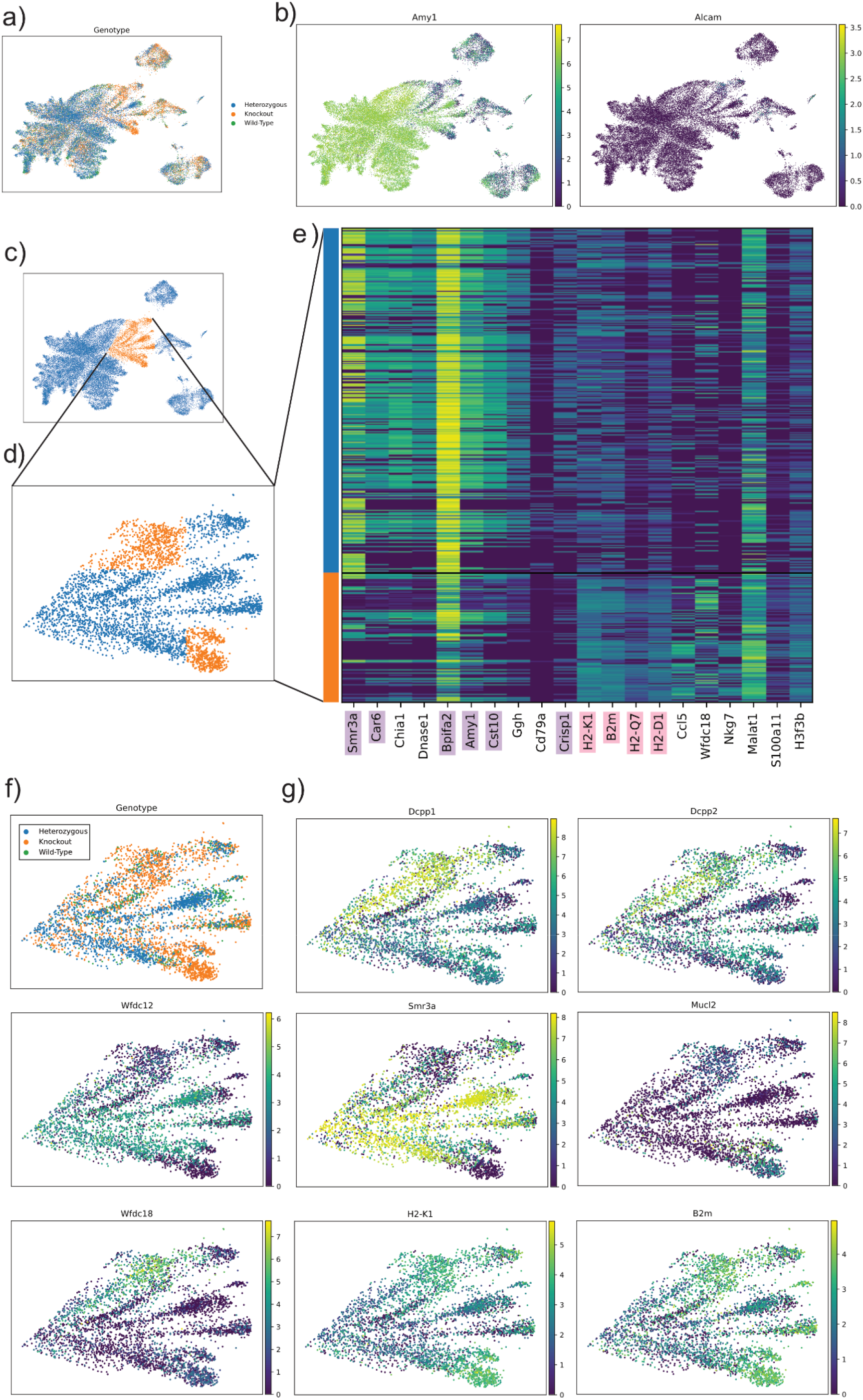
A new acinar cell type is found in the parotid glands of mice that lack Zbtb7b. (a) UMAP of single-cell gene expression of mouse parotids colored by genotype. (b) Feature plots of expression of Amy1, the alpha amylase gene, and Alcam. A minority of acinar cells have low Amy1 expression, are positive for Alcam and are enriched for knockout mice. (c) Cells are subset based on UMAP embeddings, specifically including the region of low-Amy1 acinar cells. (d)The subset cells are further divided into the Amy1-positive knockout-enriched cells (orange) and all other cells (blue). (e) Expression of the subset cells reveals lower levels of expression of proteins secreted by the salivary gland (purple) and higher levels of expression of MHC class-I genes (pink). (f) Genotypes of the subset cells. (g) Feature plots of the subset cells quantifying genes characteristic of acinar cell subtypes in adult glands. Cells present in knockout mice are positive for Dcpp1, Wfdc18 and Mucl2 and negative for Wfdc12 and Smr3a.

Cells were further separated based on UMAP embeddings into a novel class of Amy1-low largely-knockout cells alongside Amy1-high cells (Figure 5c,d). The new Amy1-low cells fail to express other parotid proteins including Smr3a and Car6, express lower levels of Bpifa2, and instead express higher levels of gene products that are part of the MHC Class I complex, including B2m, H2-K1 and other H2 genes (Figure 5e). Other markers of this subtype include the Dcpp class of proteins and the mucin-like protein Mucl2 (Figure 5g). The Dcpp1+Wfdc18+ pattern of expression loosely overlaps the pattern of expression of the adult submandibular-specific cell type called “Cluster 11.” This new cell type also shares an increased level of expression of MHC-I with the cell type “Cluster 8” in the submandibular glands of knockout mice.

Previously-published neonate parotid epithelial cells show a population positive for genes similar to the ductal-adjacent Set 1 in the developing submandibular, including Cldn3 and H2-K1 (Figure 6a, Supplementary Figure 5a). Parotid glands from knockout mice were superficially distinct at the histological level from those of heterozygotes and wild-type mice. When glands were stained for alpha amylase and Alcam, patches of high-density acinar cells were visible in homozygous mice that lacked alpha amylase and expressed Alcam (Figure 6b). These cells were not present in all sections of the glands and represented a minority of the acinar area when present.

**Figure 6.**
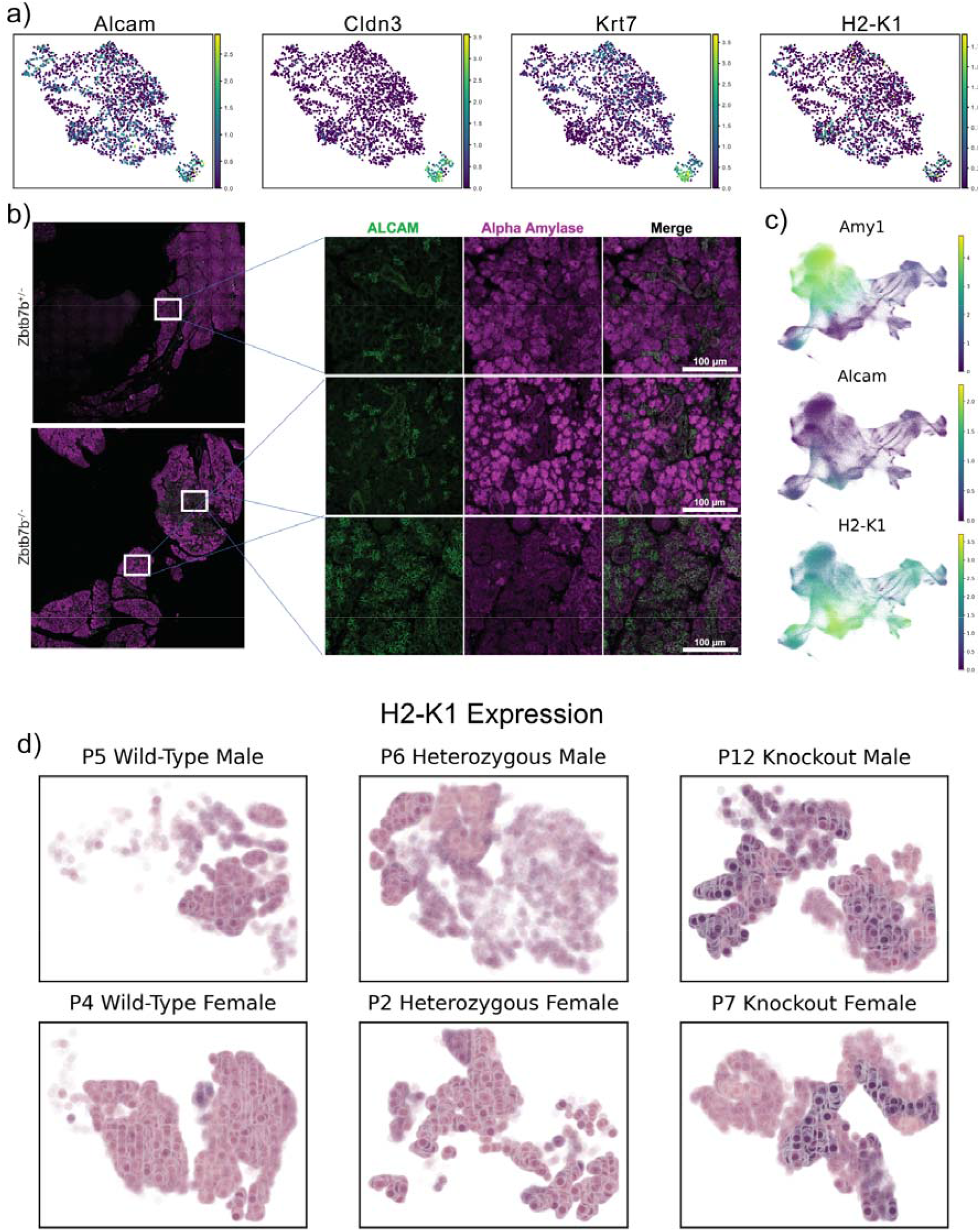
The new population of acinar cells in homozygous knockout mice resembles a H2-K1+ neonatal population lost in adulthood. (a) UMAP feature plots of single-cell RNA-seq data from neonatal mouse parotid epithelium. (b) Immunohistochemistry images from parotid glands of mice heterozygous and homozygous for Zbtb7b knockout. Heterozygous mice produce alpha amylase in acini and low levels of Alcam around ducts. Homozygous mouse parotids have multiple kinds of acini, including acini that resemble wild-type mouse glands and acini that produce no amylase and are highly Alcam positive. (c) UMAPs of Xenium data from knockout and wild-type adult parotid glands. An Alcam+H2-K1+ population of acini can be distinguished, specific to homozygous knockout mice. (d) Visualization of H2-K1 expression in the adult glands across sex and genotype. Knockout mice have glandular compartments populated by H2-K1+ acinar cells.

Acinar, ductal and endothelial cell populations in wild-type and knockout mouse parotids were distinguishable using Xenium data, and acinar populations from homozygous knockout mice could be distinguished from all cell types in heterozygous and wild-type mice (Supplementary Figure 5b,c). Wild-type and heterozygous mice had Amy1+ parotid acini, while homozygous knockout mice had both Amy1+ acini as well as a population Amy-Alcam+H2-K1+ acini (Figure 6c, Supplementary Figure 5d). These H2-K1+ acini form a well-defined and spatially segregated subset of the glands of both male and female knockout mice (Figure 6d).

## Discussion

ZBTB7B/Zbtb7b plays a role in regulating other transcription factors that makes it critical to salivary gland function. Even partial loss of this transcription factor in mice leads to a loss of salivary gland flow and complete loss of the transcription factor results in shifts in the cell populations in the submandibular gland and the emergence or retention of a neonate-like acinar cell population in the parotid. That glandular dysfunction did not accompany acinar cell loss but instead that homozygous mice developed a new, minority acinar cell population was unexpected. Significantly, these new cells in the gland have consistent and high expression of MHC class I genes involved in the innate immune response in both the parotid and submandibular glands of knockout mice. Complicating the implication of these new cells in the submandibular gland is the discrepancy between adult and developing submandibular glands.

Using a hidden Markov model (HMM) to segment the genome into different chromatin states or regulatory regions is an established approach, with the original model using a Gaussian emission distribution on ChIP-seq data(47). HMMs limited to ChIP-seq data have become increasingly sophisticated, making use of multiple hierarchical levels of hidden state(48) or multiple biological conditions(49, 50). Unfortunately, HMMs built on chromatin accessibility data are limited computationally to small genomes(51) or a low number of hidden states(52). Biologists using HMMs are increasingly opting for flexibility in emissions distribution to match the distribution of genomic data(53). Most commonly, this distribution is a negative binomial. This paper improves on previous approaches by modeling chromatin state as a set of Poisson emissions that sum to a negative binomial. The maximum likelihood estimator of a Poisson distribution is its mean, keeping expectation maximization simple. Many HMMs were trained on 4Mbp windows around genes of interest, parallelizing the problem, and HMM states were clustered on expected emissions to provide coherence across models.

There is no disease corresponding to a lack of ZBTB7B and humans with dysfunctional ZBTB7B are not known to exist. ZBTB7B is known as a silencer(54), consistent with the negative correlation between ZBTB7B binding sites and gene expression found here. However, ZBTB7B binding sites were characteristic of open chromatin regions associated with gene expression in AQP5 and CLDN3. Expression of ZBTB7B was correlated both positively and negatively with salivary- and non-salivary-specific genes, suggesting ZBTB7B may function as a silencer or activator depending on the genomic context.

One limitation of the study is the identity of the neonate-like parotid population found in knockout mouse glands. Despite differences in gene expression, cells enriched in knockout submandibular glands (“Cluster 8”) were recognizable as a ductal species. The new acinar cells expressed both ductal markers (Krt8, high Epcam) as well as acinar markers (Etv1, Dcpp2, high Aqp5) and histologically resembled ducts. They may represent ducts with aberrant gene expression, a population that has not differentiated into either acini or ducts, or a cell type not analogous to one found in normal development.

Another limitation of the study is the lack of orthology between the effects of ZBTB7B and Zbtb7b on the innate immune systems of humans and mice, respectively. Absence of Zbtb7b leads to both new or higher proportions of cell populations that express higher levels of MHC Class I genes, and a higher level of expression of those genes across the gland. However, in humans, B2M and CD74 correlate with ZBTB7B in RNA-seq from healthy salivary glands.

Openness in CD74’s hidden states all correlate with expression, no matter their span or the presence or absence of ZBTB7B binding sites, and a HMM cannot be built on the highly variable HLA region.

Despite these limitations, this study establishes the central role of ZBTB7B in salivary gland function in mice and humans. ZBTB7B is found in ducts, acini and resident immune cells and modifies their interactions and gene expression pattern, and its loss decreases salivary function. This work informs future reprogramming using master regulators such as ZBTB7B to restore function to dysfunctional salivary glands, to generate salivary organoids from iPSCs and to sustain healthy networks of gene expression during treatments stressful to salivary glands.

---

**Supplementary Figure 1.**
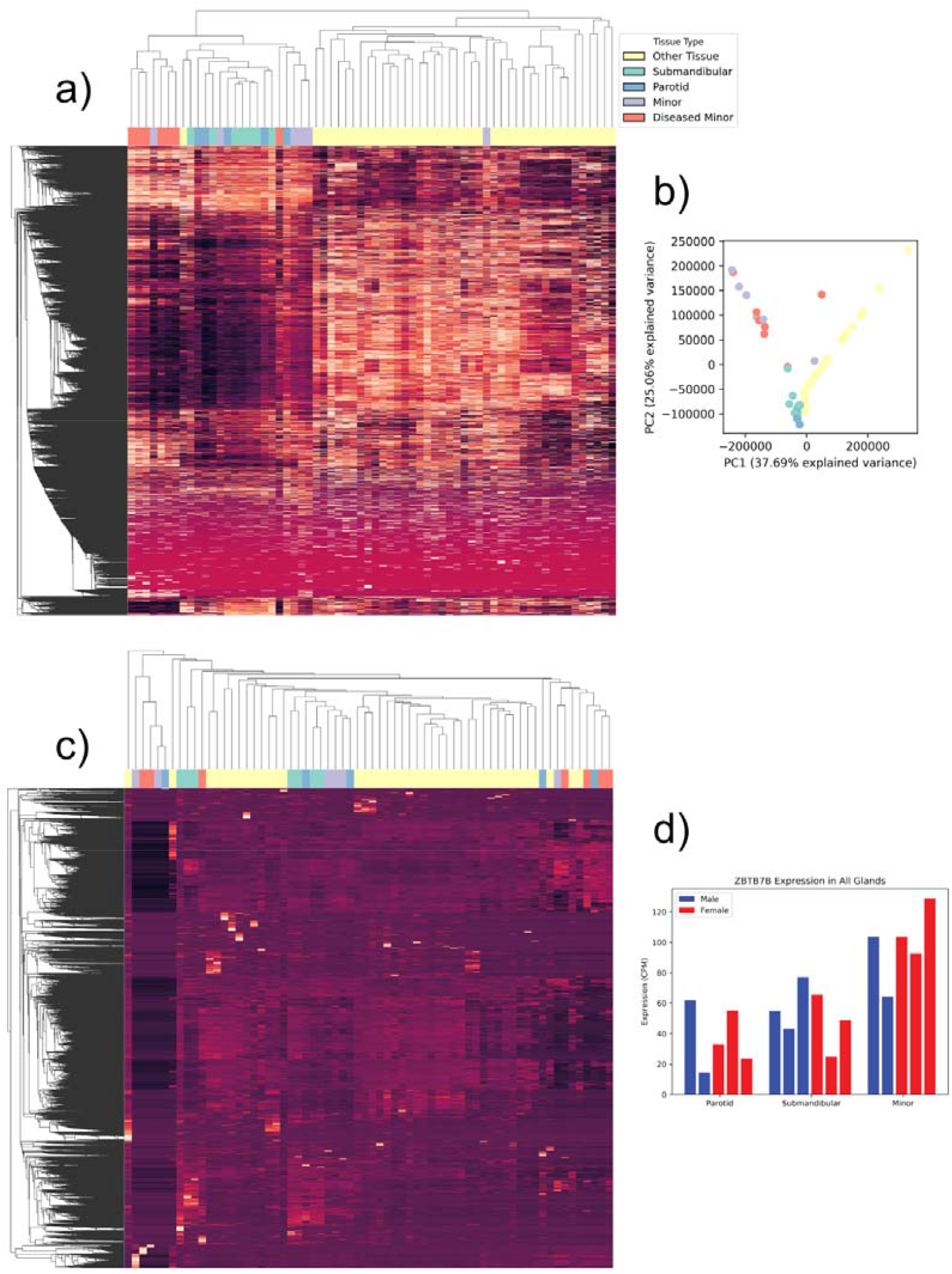
Salivary glands segregate from other tissues on gene expression but not on patterns of chromatin openness around transcription start sites (TSSs). (a) Heatmap of gene expression in salivary glands and other tissues. (b) PCA plot of tissues decomposed from gene expression data. (c) Heatmap of chromatin accessibility in 10kb windows around TSSs. (d) Expression of ZBTB7B in major and minor salivary glands.

**Supplementary Figure 2.**
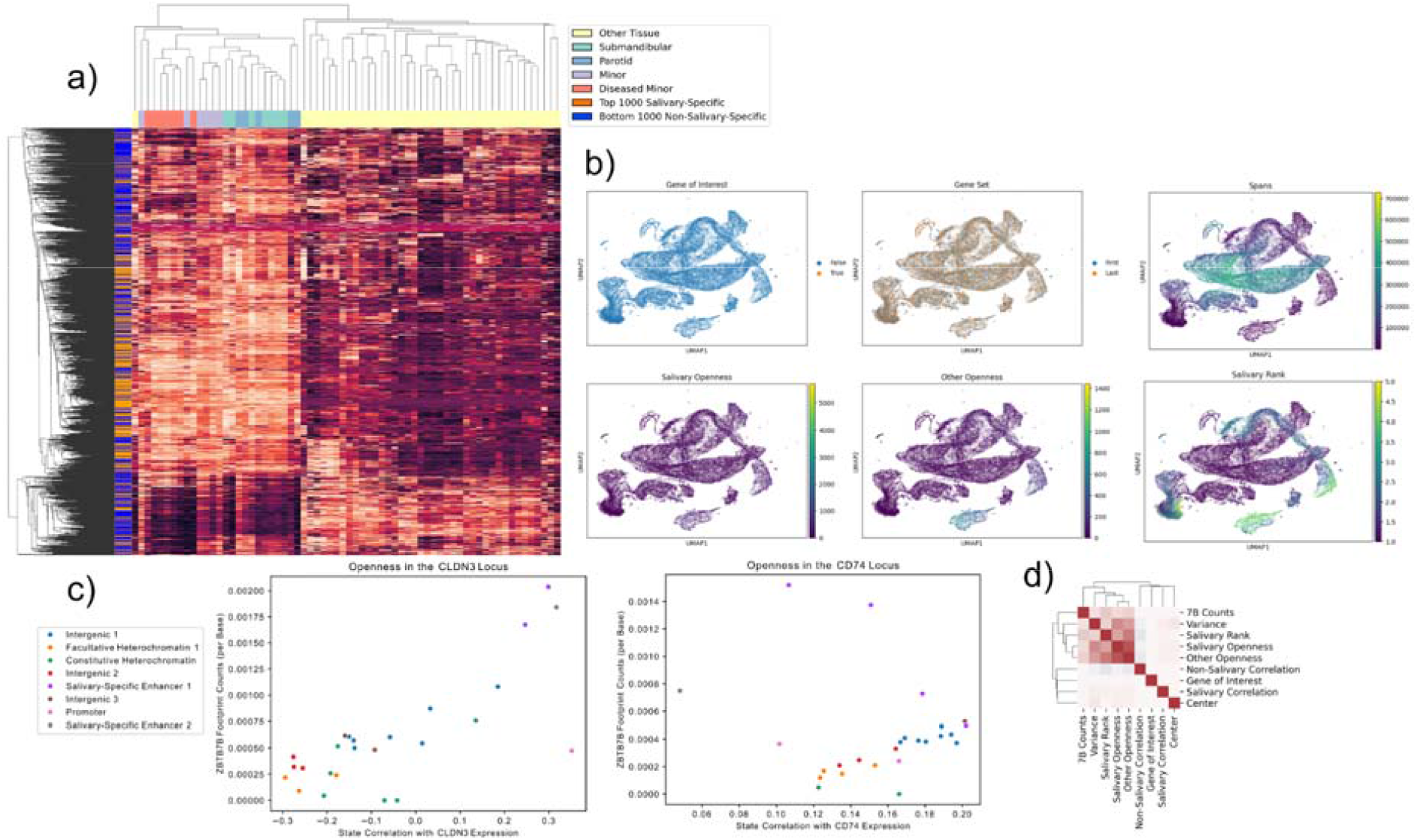
HMM states built on salivary- and non-salivary chromatin accessibility annotate genomic regions as salivary-specific and non-salivary specific. (a) Heatmap of the top 1000 salivary-specific and bottom 1000 non-salivary-specific genes. (b) Features of interest plotted in UMAP embeddings for HMM states. “Spans” is the total base pair count of a given feature within the 4Mbp window. (c) State correlations with openness and ZBTB7B site counts in CLDN3, a gene associated with salivary function, and CD74, part of the immune interface in salivary epithelium. (d) Correlations between features of interest in HMM states.

**Supplementary Figure 3.**
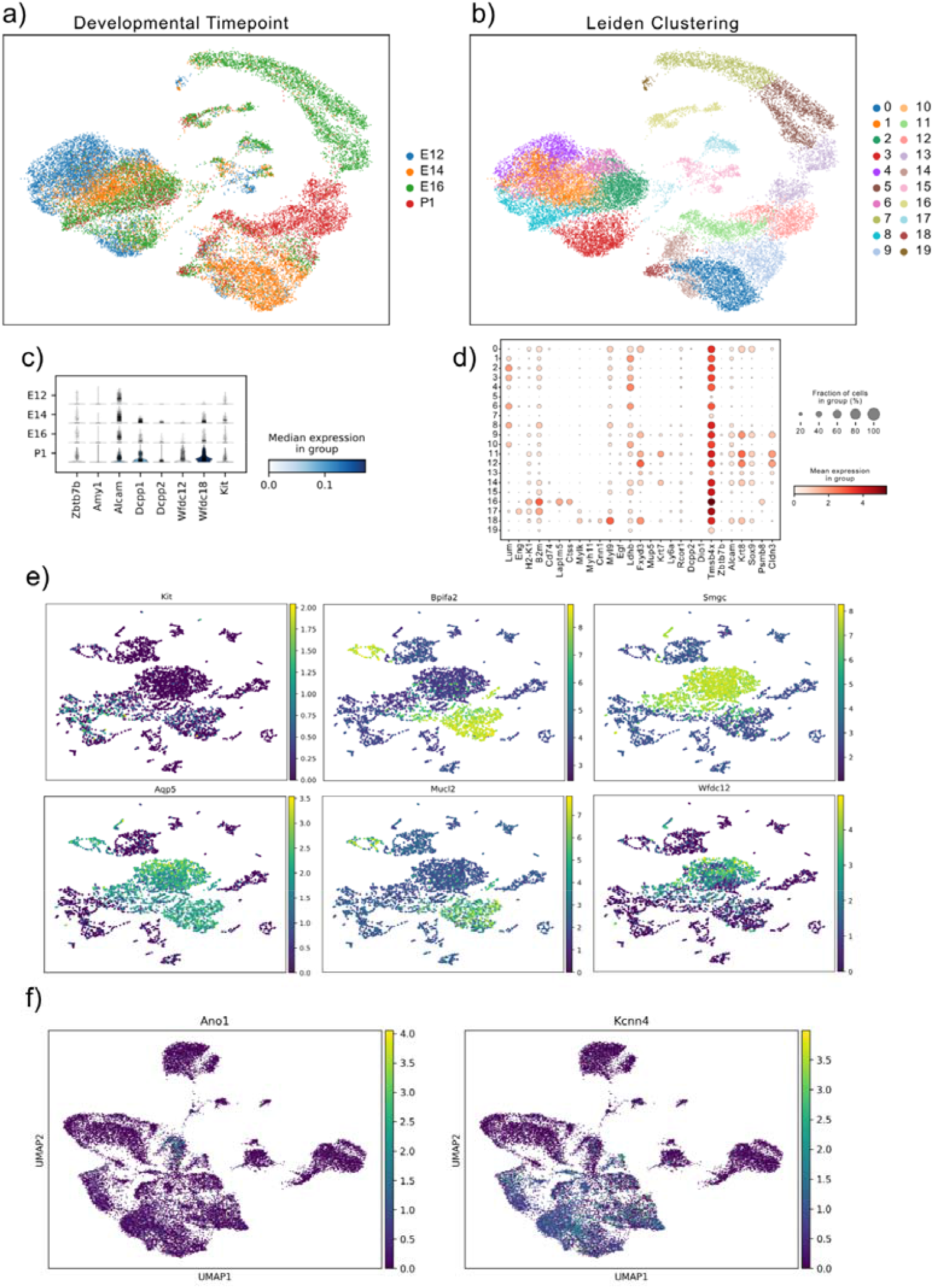
Markers of salivary function increase with developmental time alongside Zbtb7b in fetal mouse submandibular. (a) UMAP of multiple fetal mouse submandibular developmental timepoints. (b) UMAP of multiple timepoints, clustered via Leiden. (c)Changes in Zbtb7b-associated genes across multiple developmental timepoints in the fetal mouse submandibular. (d) Expression of key lineage markers and Zbtb7b-associated genes across Leiden clusters. (e) Expression of key salivary markers in the P1 sets. (f) Expression of key salivary functional genes in adult submandibular glands.

**Supplementary Figure 4.**
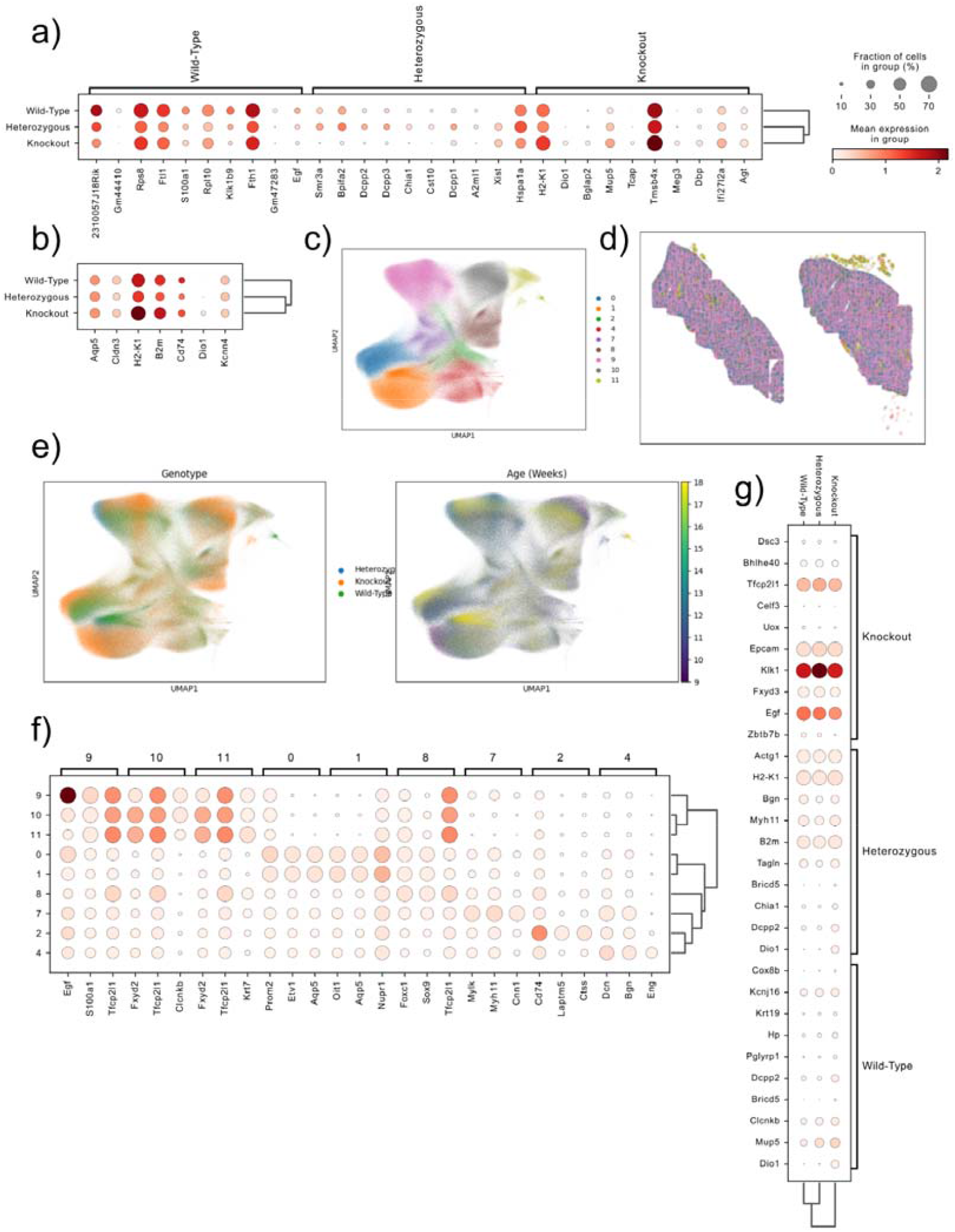
Submandibular glands of knockout and heterozygous mice have changes in expression of genes associated with salivary function and acinar identity. (a) Expression of genes associated with genotype in submandibular scRNA-seq. (b) Expression of Zbtb7b-associated genes in submandibular scRNA-seq. (c) Leiden clustering of submandibular Xenium data. (d) Visualization of Leiden clusters in physical space on the Xenium slide. (e) Genotype and age of submandibular Xenium samples, embedded in UMAP space. (f) Expression of cluster-specific gene markers in Xenium from submandibular. (g) Expression of genotype-specific genes in Xenium from submandibular.

**Supplementary Figure 5.**
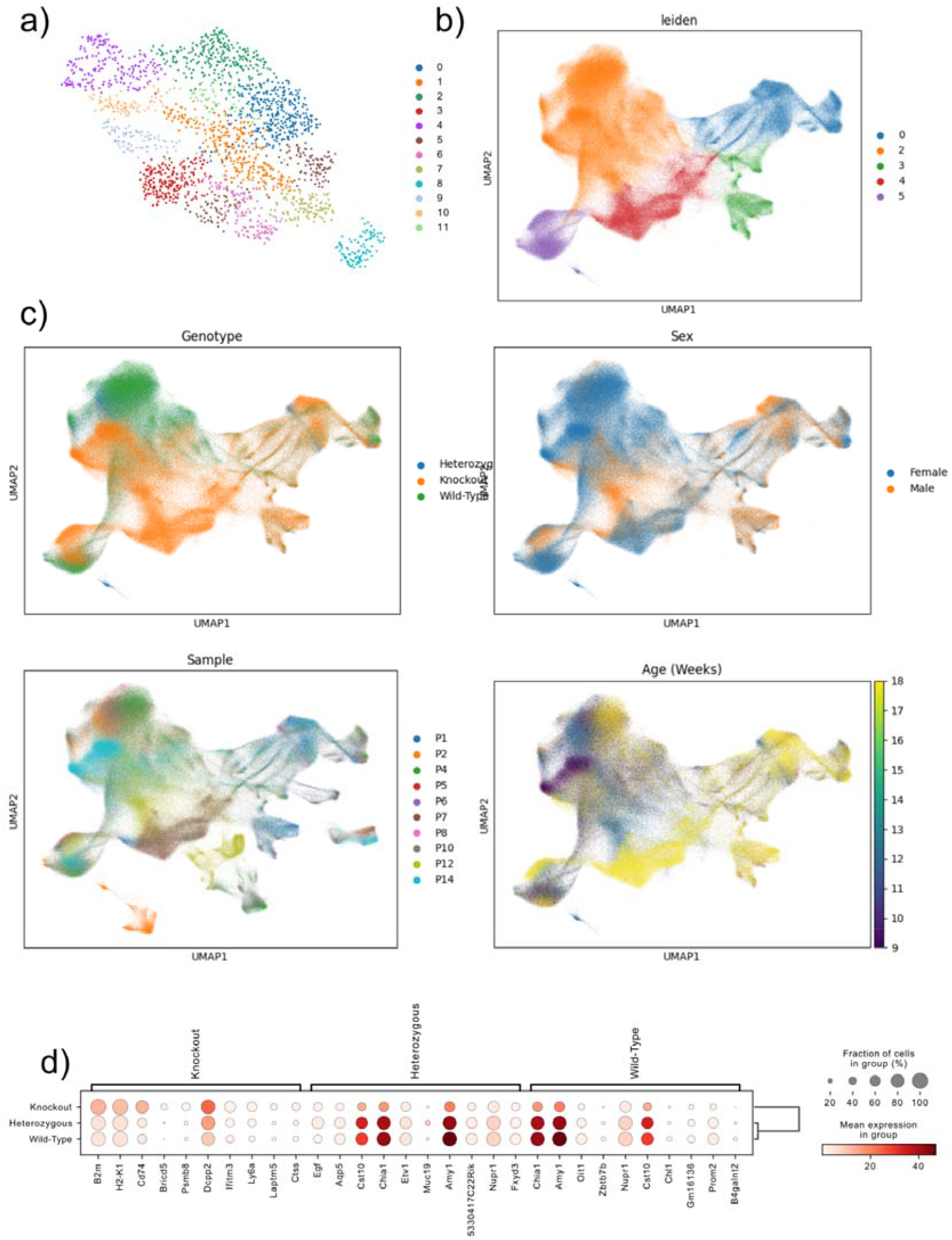
Acinar cells unique to knockout mice are found in both sexes and at multiple ages. (a) UMAP of Leiden clusters in newborn mouse parotid. (b) UMAP of Leiden clustering in Xenium of mouse adult parotid. (c) Age, sex, genotype and sample name for Xenium in mouse parotid. (d) Genotype-specific expression in Xenium from mouse parotid.

## Bibliography

1. Bolcun-Filas, E., Bannister, L.A., Barash, A., Schimenti, K.J., Hartford, S.A., Eppig, J.J., Handel, M.A., Shen, L. and Schimenti, J.C. (2011) A-MYB (MYBL1) transcription factor is a master regulator of male meiosis. Dev. Camb. Engl., 138, 3319–3330.

2. Debnath, P., Huirem, R.S., Dutta, P. and Palchaudhuri, S. (2021) Epithelial–mesenchymal transition and its transcription factors. Biosci. Rep., 42, BSR20211754.

3. Sinha, U.K. and Lin, D.-C. (2023) A novel role of master regulator transcription factor in anti-tumor immunity and cancer immunotherapy. Theranostics, 13, 1823–1825.

4. Whyte, W.A., Orlando, D.A., Hnisz, D., Abraham, B.J., Lin, C.Y., Kagey, M.H., Rahl, P.B., Lee, T.I. and Young, R.A. (2013) Master Transcription Factors and Mediator Establish Super-Enhancers at Key Cell Identity Genes. Cell, 153, 307–319.

5. Zhou, C., Wang, M., Zhang, C. and Zhang, Y. (2025) The transcription factor GABPA is a master regulator of naive pluripotency. Nat. Cell Biol., 27, 48–58.

6. Emmerson, E., May, A.J., Nathan, S., Cruz-Pacheco, N., Lizama, C.O., Maliskova, L., Zovein, A.C., Shen, Y., Muench, M.O. and Knox, S.M. SOX2 regulates acinar cell development in the salivary gland. eLife, 6, e26620.

7. Vacchio, M.S., Ciucci, T., Gao, Y., Watanabe, M., Balmaceno-Criss, M., McGinty, M.T., Huang, A., Xiao, Q., McConkey, C., Zhao, Y., et al. (2019) A Thpok-Directed Transcriptional Circuitry Promotes Bcl6 and Maf Expression to Orchestrate T Follicular Helper Differentiation. Immunity, 51, 465–478.e6.

8. Vacchio, M.S., Wang, L., Bouladoux, N., Carpenter, A.C., Xiong, Y., Williams, L.C., Wohlfert, E., Song, K.-D., Belkaid, Y., Love, P.E., et al. (2014) A ThPOK-LRF transcriptional node maintains the integrity and effector potential of post-thymic CD4+ T cells. Nat. Immunol., 15, 947–956.

9. Ciucci, T., Vacchio, M.S., Gao, Y., Tomassoni Ardori, F., Candia, J., Mehta, M., Zhao, Y., Tran, B., Pepper, M., Tessarollo, L., et al. (2019) The Emergence and Functional Fitness of Memory CD4+ T Cells Require the Transcription Factor Thpok. Immunity, 50, 91–105.e4.

10. Gao, Y., Zamisch, M., Vacchio, M., Chopp, L., Ciucci, T., Paine, E.L., Lyons, G.C., Nie, J., Xiao, Q., Zvezdova, E., et al. (2022) NuRD complex recruitment to Thpok mediates CD4 + T cell lineage differentiation. Sci. Immunol., 7, eabn5917.

11. Mariani, F., Sena, P., Pedroni, M., Benatti, P., Manni, P., Di Gregorio, C., Manenti, A., Palumbo, C., De Leon, M.P. and Roncucci, L. (2013) Th Inducing POZ-Kruppel Factor (ThPOK) Is a Key Regulator of the Immune Response since the Early Steps of Colorectal Carcinogenesis. PLoS ONE, 8, e54488.

12. Kennedy, J.M., Georges, A., Bassenden, A.V., Vidal, S.M., Berghuis, A.M., Taniuchi, I., Majewski, J., Lathrop, M., Behr, M.A., Langlais, D., et al. (2020) ZBTB7B (ThPOK) Is Required for Pathogenesis of Cerebral Malaria and Protection against Pulmonary Tuberculosis. Infect. Immun., 88, e00845–19.

13. Zhao, J., Han, D.-X., Wang, C.-B. and Wang, X.-L. (2020) Zbtb7b suppresses aseptic inflammation by regulating m6A modification of IL6 mRNA. Biochem. Biophys. Res. Commun., 530, 336–341.

14. De Arcos-Jiménez, J.C., González-Hernández, L.A., Ratkovich-González, S., Sánchez-Reyes, K., Alvarez-Zavala, M., Ruiz-Briseño, M.D.R., Mosqueda-Gómez, J.L., Avila-Rios, S., Ramos-Solano, M. and Andrade-Villanueva, J.F. (2021) ZBTB gene expression in HIV patients: a possible new molecular mechanism of viral control. Arch. Virol., 166, 167–178.

15. Basu, J., Olsson, A., Ferchen, K., Titerina, E.K., Chetal, K., Nicolas, E., Czyzewicz, P., Levchenko, D., Ge, L., Hua, X., et al. (2023) ThPOK is a critical multifaceted regulator of myeloid lineage development. Nat. Immunol., 24, 1295–1307.

16. Zou, W., Izawa, T., Rohatgi, N., Zou, S.Y., Li, Y. and Teitelbaum, S.L. (2022) THPOK Inhibits Osteoclast Formation Via NFATC1 Transcription and Function. JBMR Plus, 6, e10613.

17. Ghayor, C. (2000) Regulation of human COL2A1 gene expression in chondrocytes: identification of C-KROX responsive elements and modulation by phenotype alteration. J. Biol. Chem., 10.1074/jbc.M002139200.

18. Kypriotou, M., Beauchef, G., Chadjichristos, C., Widom, R., Renard, E., Jimenez, S.A., Korn, J., Maquart, F.-X., Oddos, T., Von Stetten, O., et al. (2007) Human Collagen Krox Up-regulates Type I Collagen Expression in Normal and Scleroderma Fibroblasts through Interaction with Sp1 and Sp3 Transcription Factors. J. Biol. Chem., 282, 32000–32014.

19. Renard, E., Chadjichristos, C., Kypriotou, M., Beauchef, G., Bordat, P., Dompmartin, A., Widom, R.L., Boumediene, K., Pujol, J. and Galéra, P. (2008) Chondroitin sulphate decreases collagen synthesis in normal and scleroderma fibroblasts through a SmadLindependent TGFLβ pathway – implication of CLKrox and Sp1. J. Cell. Mol. Med., 12, 2836–2847.

20. Zhang, R., Ma, H., Gao, Y., Wu, Y., Qiao, Y., Geng, A., Cai, C., Han, Y., Zeng, Y.A., Liu, X., et al. (2018) Th-POK regulates mammary gland lactation through mTOR-SREBP pathway. PLOS Genet., 14, e1007211.

21. Arcuschin, C.D., Kahrizi, K., Sayaman, R.W., DiBenedetto, C., Salaberry, P.J., Pirker, R., Shen, Y., Zakraoui, O., Schwarzer, C., Scapozza, A., et al. (2025) Super-enhancer profiling reveals ThPOK/ZBTB7B, a CD4+ cell lineage commitment factor, as a master regulator that restricts breast cancer cells to a luminal non-migratory phenotype. Cell. Mol. Life Sci., 82, 397.

22. Jeong, Y., Oh, A.-R., Jung, Y.H., Jung, K.H., Lee, S., Carrer, M., Lee, S.B., Valenti, L., Pajvani, U.B. and Kim, K. (2025) Hepatocyte KCTD17-mediated SERPINA3 inhibition determines liver fibrosis in metabolic dysfunction-associated steatohepatitis. Exp. Mol. Med., 57, 1673–1685.

23. Han, Y., Wu, K., Peng, X., Fu, Y., Li, W., Ma, J., Jiang, H. and Zhao, X. (2024) Zbtb7b defines a compensatory mechanism in MASLD Lrelated HCC progression by suppressing H19 Lmediated hepatic lipid deposition. Physiol. Rep., 12, e70160.

24. Zhu, Y., Wang, Q., Xie, X., Ma, C., Qiao, Y., Zhang, Y., Wu, Y., Gao, Y., Jiang, J., Liu, X., et al. (2024) ZBTB7B is a permissive regulator of hepatocellular carcinoma initiation by repressing c-Jun expression and function. Cell Death Dis., 15, 55.

25. Hong, F., Wang, X., Zhong, N., Zhang, Z., Lin, S., Zhang, M., Li, H., Liu, Y., Wang, Y., Zhao, L., et al. (2025) The critical role of BMP signaling in gastric epithelial cell differentiation revealed by organoids. Cell Regen., 14, 18.

26. Chen, Y., Jiang, L., Xia, L., Zhang, G. and Chen, L. (2022) ThPOK inhibits the immune escape of gastric cancer cells by inducing STPG1 to inactivate the ERK pathway. BMC Immunol., 23, 16.

27. Cui, K., Yao, S., Liu, B., Sun, S., Gong, L., Li, Q., Fei, B. and Huang, Z. (2022) A novel high-risk subpopulation identified by CTSL and ZBTB7B in gastric cancer. Br. J. Cancer, 127, 1450–1460.

28. Liu, F.-P., Huang, T., Yang, H.-Y., Zha, J.-H., Xu, C., Qiu, C.-P., Wang, X., Lu, Z. and Wang, T. (2026) Inhibition of ZBTB7B-mediated ADPGK transcription by NEDD4 impedes glycolysis and progression of lung adenocarcinoma. Oncogenesis, 15, 14.

29. Liu, L., Gao, Z., Wang, Q., Zhao, X., Zhang, F., Zhu, Y., Sun, W., Huang, Y., Wang, Z., Yan, S., et al. (2026) TRIM29 promotes pancreatic cancer MVI via IκBα K48-ubiquitination and NF-κB activation in CXCL5+ epithelial cells. Mol. Cancer, 25, 145.

30. Zhu, L., Feng, H., Jin, S., Tan, M., Gao, S., Zhuang, H., Hu, Z., Wang, H., Song, Z. and Lin, B. (2017) High expressions of BCL6 and Lewis y antigen are correlated with high tumor burden and poor prognosis in epithelial ovarian cancer. Tumor Biol., 39, 101042831771165.

31. Heide, T., Maurer, A., Eipel, M., Knoll, K., Geelvink, M., Veeck, J., Knuechel, R., Van Essen, J., Stoehr, R., Hartmann, A., et al. (2019) Multiregion human bladder cancer sequencing reveals tumour evolution, bladder cancer phenotypes and implications for targeted therapy. J. Pathol., 248, 230–242.

32. Lee, W., Lavery, L., Rousseaux, M.W.C., Rutledge, E.B., Jang, Y., Wan, Y., Wu, S., Kim, W., AlLRamahi, I., Rath, S., et al. (2021) Dual targeting of brain regionLspecific kinases potentiates neurological rescue in Spinocerebellar ataxia type 1. EMBO J., 40, EMBJ2020106106.

33. Lin, S., Wang, C., Li, Z. and Qiu, X. (2023) Distinct H3K27me3 and H3K27ac Modifications in Neural Tube Defects Induced by Benzo[a]pyrene. Brain Sci., 13, 334.

34. Li, S., Mi, L., Yu, L., Yu, Q., Liu, T., Wang, G.-X., Zhao, X.-Y., Wu, J. and Lin, J.D. (2017) Zbtb7b engages the long noncoding RNA Blnc1 to drive brown and beige fat development and thermogenesis. Proc. Natl. Acad. Sci., 114.

35. Zhao, X.-Y., Zhou, L., Chen, Z., Ji, Y., Peng, X., Qi, L., Li, S. and Lin, J.D. (2020) The obesity-induced adipokine sST2 exacerbates adipose Treg and ILC2 depletion and promotes insulin resistance. Sci. Adv., 6, eaay6191.

36. Turcot, V., Lu, Y., Highland, H.M., Schurmann, C., Justice, A.E., Fine, R.S., Bradfield, J.P., Esko, T., Giri, A., Graff, M., et al. (2018) Protein-altering variants associated with body mass index implicate pathways that control energy intake and expenditure in obesity. Nat. Genet., 50, 26–41.

37. Cao, H., Huang, H. and Tang, H. (2021) Zbtb7a and Zbtb7b: Opening naïve loci to reprogram ESCs. Biosci. Trends, 15, 58–60.

38. Buenrostro, J.D., Wu, B., Chang, H.Y. and Greenleaf, W.J. (2015) ATACLseq: A Method for Assaying Chromatin Accessibility GenomeLWide. Curr. Protoc. Mol. Biol., 109.

39. Pranzatelli, T.J.F., Michael, D.G. and Chiorini, J.A. (2018) ATAC2GRN: optimized ATAC-seq and DNase1-seq pipelines for rapid and accurate genome regulatory network inference. BMC Genomics, 19, 563.

40. Patro, R., Duggal, G., Love, M.I., Irizarry, R.A. and Kingsford, C. (2017) Salmon provides fast and bias-aware quantification of transcript expression. Nat. Methods, 14, 417–419.

41. Srivastava, A., Malik, L., Sarkar, H., Zakeri, M., Almodaresi, F., Soneson, C., Love, M.I., Kingsford, C. and Patro, R. (2020) Alignment and mapping methodology influence transcript abundance estimation. Genome Biol., 21, 239.

42. Weirauch, M.T., Yang, A., Albu, M., Cote, A.G., Montenegro-Montero, A., Drewe, P., Najafabadi, H.S., Lambert, S.A., Mann, I., Cook, K., et al. (2014) Determination and Inference of Eukaryotic Transcription Factor Sequence Specificity. Cell, 158, 1431–1443.

43. Grant, C.E., Bailey, T.L. and Noble, W.S. (2011) FIMO: scanning for occurrences of a given motif. Bioinformatics, 27, 1017–1018.

44. McInnes, L., Healy, J. and Melville, J. (2020) UMAP: Uniform Manifold Approximation and Projection for Dimension Reduction.

45. V. A. Traag, L. Waltman, and N. J. van Eck (2019) From Louvain to Leiden: guaranteeing well-connected communities. Sci. Rep., 9.

46. Wolf, F.A., Angerer, P. and Theis, F.J. (2018) SCANPY: large-scale single-cell gene expression data analysis. Genome Biol., 19, 15.

47. Ernst, J. and Kellis, M. (2012) ChromHMM: automating chromatin-state discovery and characterization. Nat. Methods, 9, 215–216.

48. Marco, E., Meuleman, W., Huang, J., Glass, K., Pinello, L., Wang, J., Kellis, M. and Yuan, G.-C. (2017) Multi-scale chromatin state annotation using a hierarchical hidden Markov model. Nat. Commun., 8, 15011.

49. Baldoni, P.L., Rashid, N.U. and Ibrahim, J.G. (2022) Efficient detection and classification of epigenomic changes under multiple conditions. Biometrics, 78, 1141–1154.

50. Leone, M., Galeota, E., Masseroli, M. and Pelizzola, M. (2022) Identification, semantic annotation and comparison of combinations of functional elements in multiple biological conditions. Bioinformatics, 38, 1183–1190.

51. Mitra, S., Zhong, J., MacAlpine, D.M. and Hartemink, A.J. (2020) RoboCOP: Multivariate State Space Model Integrating Epigenomic Accessibility Data to Elucidate Genome-Wide Chromatin Occupancy. Res. Comput. Mol. Biol., 12074, 136–151.

52. Shpynov, O. and Artyomov, M.N. (2026) Accurate chromatin marks peak calling with Omnipeak. Nucleic Acids Res., 54, gkaf1454.

53. Schmitz, J.E., Aggarwal, N., Laufer, L., Walter, J., Salhab, A. and Rahmann, S. (2024) EpiSegMix: a flexible distribution hidden Markov model with duration modeling for chromatin state discovery. Bioinformatics, 40, btae178.

54. Mookerjee-Basu, J., Hua, X., Ge, L., Nicolas, E., Li, Q., Czyzewicz, P., Zhongping, D., Peri, S., FuxmanBass, J.I., Walhout, A.J.M., et al. (2019) Functional Conservation of a Developmental Switch in Mammals since the Jurassic Age. Mol. Biol. Evol., 36, 39–53.

